# Forecasting Urban Wastewater Microbiome Dynamics Using a Digital Twin Framework

**DOI:** 10.1101/2025.07.21.666059

**Authors:** Bichar Dip Shrestha Gurung, Manish Rayamajhi, Naina Maharjan, Tuyen Do, Dikshya Bhandari, Rupesh Yadav, Shiva Aryal, Etienne Z. Gnimpieba

**Author notes:** these authors contributed equally to this work (Bichar Dip Shrestha Gurung, Manish Rayamajhi).

## Abstract

Urban wastewater microbiomes are complex and temporally dynamic, offering valuable insight into community-scale microbial ecology and potential public health trends. However, existing wastewater-based studies often remain descriptive, lacking tools for predictive modeling. In this study, we introduce a digital twin framework that forecasts microbial abundance trajectories in urban wastewater using an interpretable generative model, Q-net. Trained on a 30-week longitudinal metagenomic dataset from seven wastewater treatment plants, the model captures temporal microbial dynamics with high fidelity (R^2^ > 0.97 for key taxa; R^2^ = 0.998 at the final timepoint). Beyond accurate forecasting, Q-net provides transparent model structure through conditional inference trees and enables simulation of realistic microbial trends under hypothetical scenarios. This work demonstrates the potential of digital twins to move wastewater microbiome studies from static snapshots to dynamic, predictive systems, with broad implications for environmental monitoring and microbial ecosystem modeling.

## 1 Introduction

In an era of increasing global connectivity and emergent health threats, proactive public health surveillance is paramount. Traditional clinical and syndromic surveillance methods, while critical, often face limitations in scalability, timeliness, and representativeness, especially at the population level^1^. Wastewater-based epidemiology (WBE) has rapidly emerged as a powerful, non-invasive, and cost-effective tool to circumvent these challenges, providing critical population-level insights into the circulation of pathogens, antimicrobial resistance (AMR)^2^, and other public health indicators within urban environments^3^. The utility of this approach has been notably underscored by its application in tracking the spread of SARS-CoV-2 and other infectious agents^4^.

However, the urban wastewater ecosystem is inherently complex. It is characterized by a vast diversity of microorganisms originating not only from human waste, but also from industrial discharges, environmental runoff, and crucially, established sewer infrastructure biofilms^5,6^. This inherent complexity, coupled with significant temporal and spatial variability, poses substantial challenges to comprehensively understanding and predictively leveraging wastewater microbiome data for public health. Microbial biofilms, communities of microbes attached to surfaces and encased in a self-secreted matrix, are ubiquitous in wastewater infrastructure^7^. These biofilms serve as significant, yet often overlooked, reservoirs for a diverse array of microorganisms, including pathogens and AMR-carrying bacteria, profoundly influencing water quality, treatment efficiency, and the overall dynamics of the wastewater microbiome^8^. Understanding the formation, maturation, and shedding of these biofilms is therefore crucial for a holistic comprehension of wastewater systems. While traditional metagenomic analyses of wastewater provide invaluable snapshots of microbial composition and gene content, revealing correlations with environmental factors or public health outcomes, these descriptive and correlative studies frequently fall short of providing the predictive and mechanistic understanding required for proactive public health interventions or optimized wastewater management^9,10^.

To address these critical limitations and unlock the full predictive potential of wastewater surveillance, we propose and implement a novel digital twin framework for the urban wastewater microbiome. Inspired by the digital twin paradigm in other complex biological systems^11,12^, we adopt a data-driven approach to model and forecast the temporal dynamics of the wastewater microbiome. While not a full virtual replica, our framework leverages high-resolution, longitudinal multi-omics data to emulate key microbial and functional trajectories, enabling high-fidelity prediction and actionable insights. Recent advances in generative AI and ingredient-level modeling^13^ further reinforce the viability of using hybrid machine learning and multi-modal sensing approaches in complex biological contexts. We hypothesize that by leveraging this advanced modeling paradigm, we can transcend the current descriptive limitations of WBE to achieve truly predictive and mechanistic insights into wastewater-borne public health threats. By modeling the real-time behavior of microbial taxa and environmental interactions, a digital twin enables predictive intelligence and source attribution, moving beyond static correlation toward causal understanding and actionable forecasting.

In this study, we leverage a longitudinal dataset from seven distinct wastewater treatment plants, encompassing 30 weeks of sampling and rich multi-omics data, including both shotgun sequencing and MAGs. Our primary objective is to infer a digital twin of these wastewater microbiomes that enables accurate forecasting by developing a robust model to predict the coupled dynamics of microbial species over an extended period. Additionally, we aim to achieve enhanced source attribution by reliably distinguishing and quantifying the contributions of human and environmental sources to the overall microbial community.

## 2 Results

While Budapest exhibits periodic spikes in *Psychrobacter* abundance. The microbial communities in Copenhagen sites show more balanced distributions, though transient blooms, such as a peak in *Cloacibacterium* in RD, are evident. These trends underscore the dynamic and site-specific nature of wastewater microbial ecosystems, potentially influenced by localized environmental conditions, operational differences in treatment infrastructure, or biofilm-associated shedding events.

### 2.1 Digital Twin Construction and Forecasting of Wastewater Microbiome Trajectories

To evaluate the predictive capabilities of Q-net on longitudinal sewage metagenomic data, we trained the model using observations between weeks 60–90, an interval with consistent sampling coverage across sites (see Figure 9).

Figure 2 summarizes population-level forecasts for selected taxa. Each panel shows the observed (red) versus Q-net-predicted (blue) relative abundance trajectories across the 30-week window. The shaded region around the forecast represents the ensemble confidence interval. Q-net captured both global and taxon-specific dynamics with high fidelity, achieving *R*^2^ values above 0.97 for all plotted taxa, including *Acidovorax, Propionibacterium, Aeromonas, Pseudomonas*, and *Psychrobacter*.

**Figure 1.**
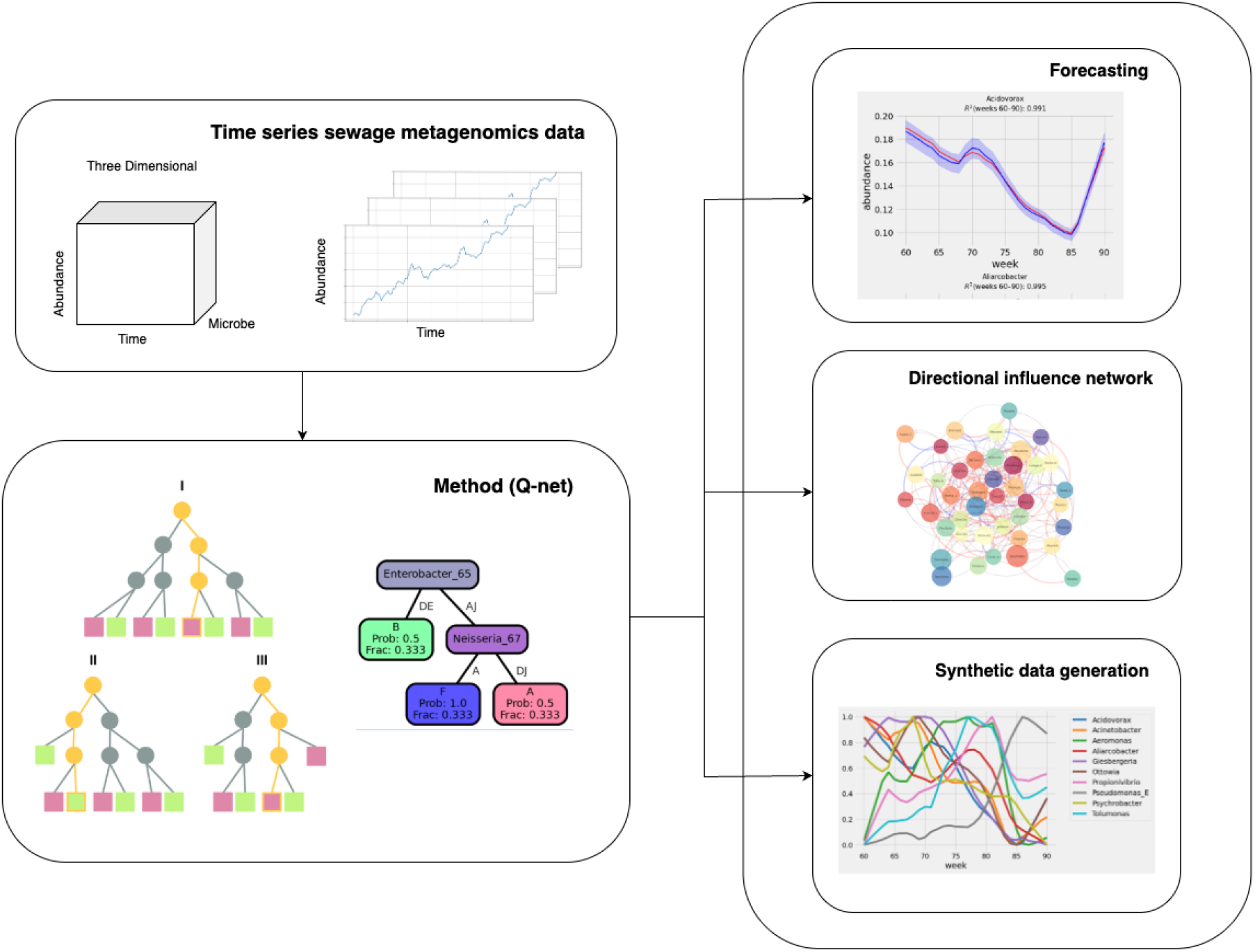
Overview of the Model Framework. Starting from time-series sewage metagenomic data, the model learns interpretable probabilistic decision ensembles to model taxon abundance trajectories. The framework supports (i) temporal forecasting, (ii) inference of directional influence networks, and (iii) simulation of phantom or synthetic subjects under data-sparse conditions.

**Figure 2.**
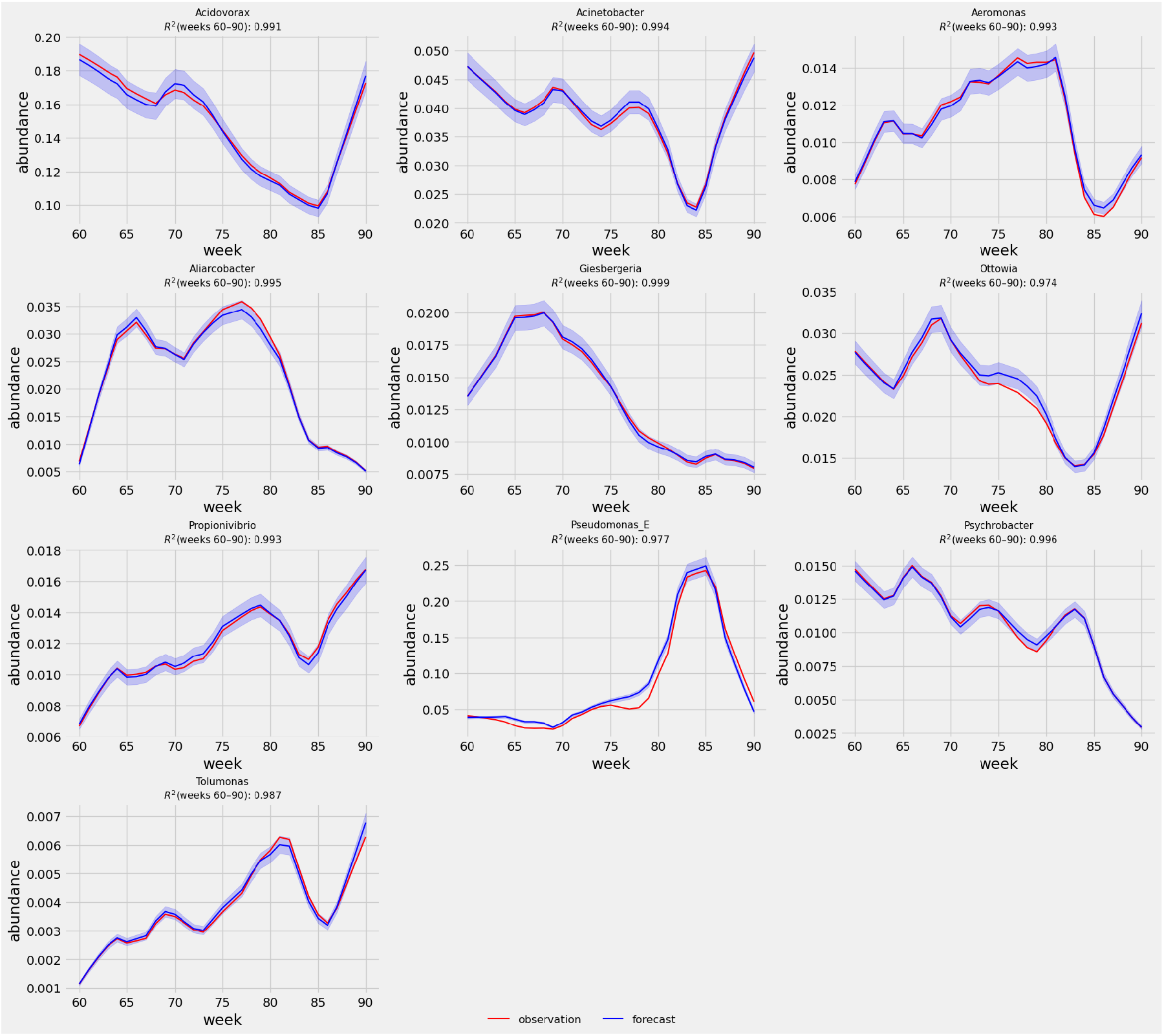
Digital Twin Forecasting Performance using Q-net. Forecasting results for selected taxa. Red lines indicate observed relative abundances; blue lines show Q-net forecasts with shaded regions denoting ensemble variability. Q-net achieved high predictive accuracy (*R*^2^ *>* 0.97) across all shown taxa during weeks 60–90.

To further validate Q-net’s predictive accuracy, we evaluated performance at the final forecast horizon (Week 90) across all genera in the dataset. As shown in Figure 3, the predicted relative abundances closely matched observed values across a wide dynamic range, yielding an *R*^2^ = 0.998. A zoomed-in inset emphasizes Q-net’s fidelity even for low-abundance taxa. These results demonstrate the digital twin’s precision in capturing endpoint community structure.

**Figure 3.**
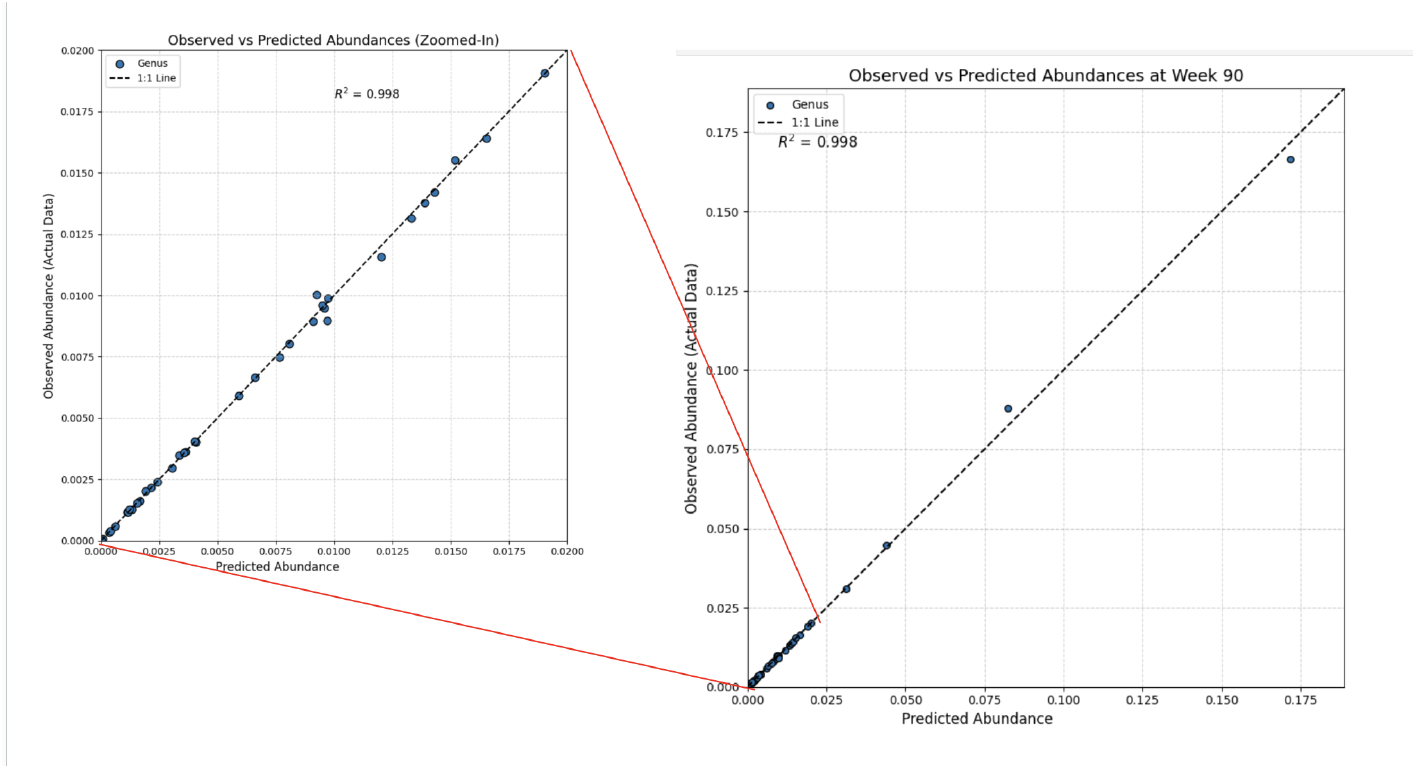
Observed vs. Predicted Abundances at Week 90. Scatter plot of Q-net-predicted vs. observed relative abundances for all genera at the final forecast timepoint (Week 90). The model achieved a near-perfect *R*^2^ = 0.998. The inset shows a zoomed-in view of low-abundance taxa, confirming accuracy across the entire distribution.

In addition to accurate forecasting, Q-net offers a high degree of interpretability by modeling each microbial taxon using a conditional inference tree (CIT). These trees estimate the conditional distribution of a target taxon at a given timepoint, conditioned on the quantized abundance states of other taxa across past timepoints. Each CIT is constructed using a statistically principled, permutation-based splitting criterion that avoids the variable selection bias common in standard decision trees^14^.

Figure 4 presents a representative CIT from the Q-net ensemble trained on longitudinal sewage metagenomic data. The tree models the abundance state of *Enterobacter_65*, revealing that the state of *Neisseria_67* and other taxa at specific prior timepoints are strong predictors of future dynamics. Each node in the tree shows the predicted class distribution and the fraction of training samples passing through that node.

**Figure 4.**
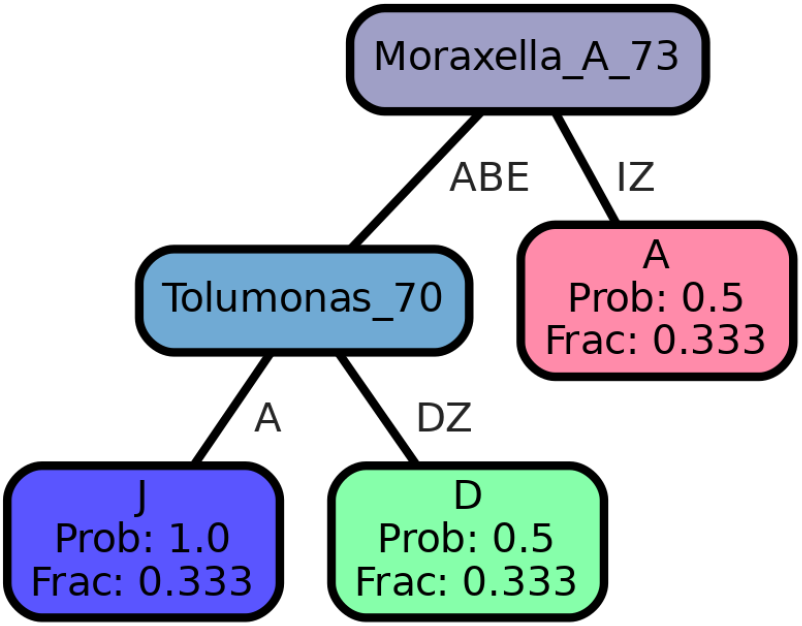
Example conditional inference tree from Q-net. This tree models the abundance state of *Moraxella_A_65* based on past abundance states of other taxa. Nodes indicate predicted bin probabilities and fraction of training samples.

These tree structures not only support accurate forecasting, but also help identify latent ecological dependencies, temporal lag effects, and taxon–taxon influence pathways. This interpretability advantage distinguishes Q-net from more opaque modeling approaches such as neural networks or transformers^15^.

### 2.2 Visualization of the Microbial Regulatory Network

Building on the LOMAR coefficients estimated via Q-net’s marginalization procedures, we visualized the directed influence network among microbial taxa. Each edge represents a hypothesized regulatory relationship, derived by computing the marginal change in the abundance of a target taxon due to perturbation in a source taxon.

As shown in Figure 5, the network reveals a dense web of directional dependencies, with certain taxa (e.g., *Tolumonas, Enterobacter, Rubrivivax*) exhibiting high centrality, potentially indicating their regulatory significance in shaping microbial community dynamics.

**Figure 5.**
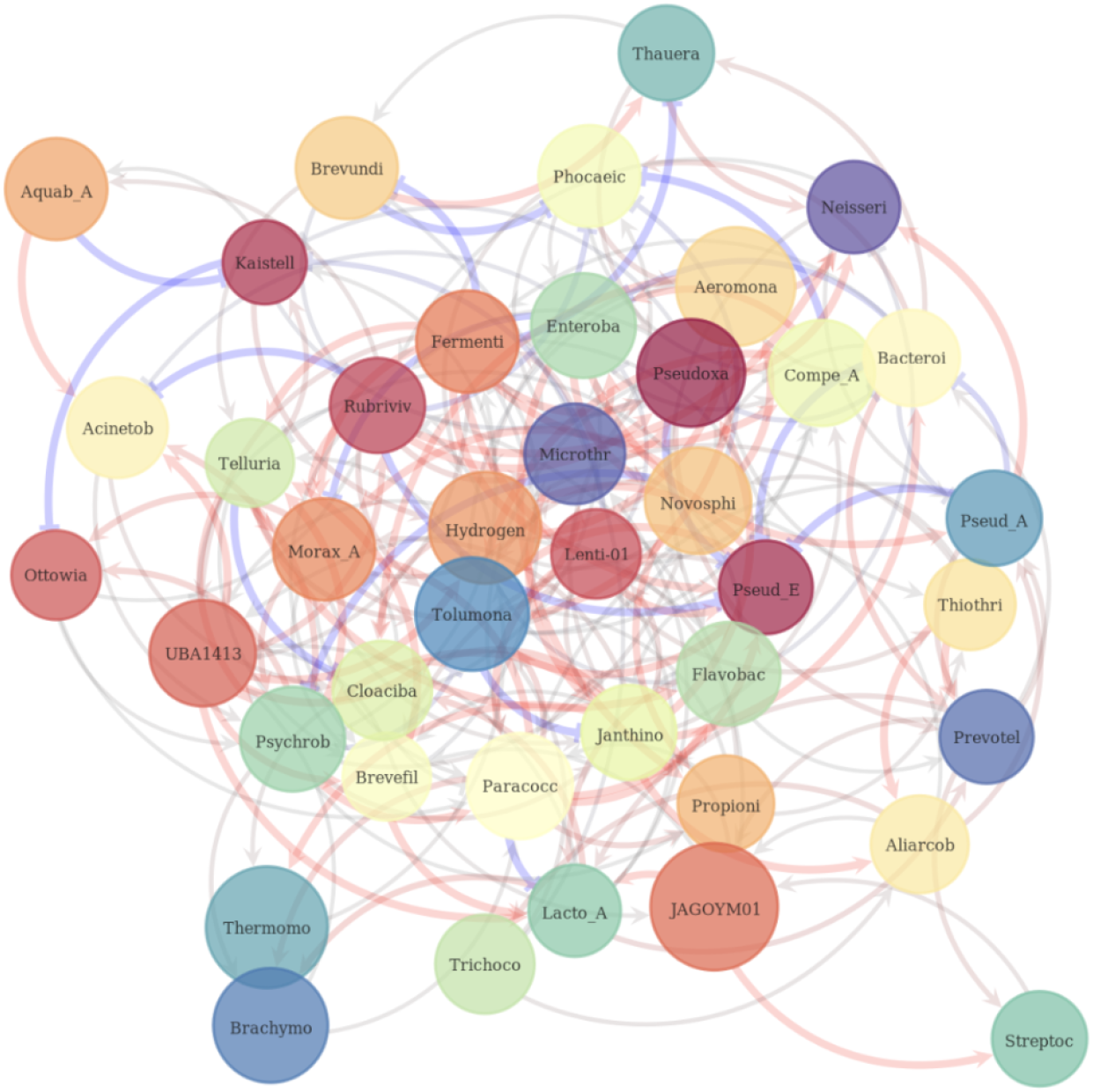
Directional influence network inferred via Q-net-LOMAR. Nodes represent microbial genera. Directed edges denote marginal influence relationships inferred from Q-net conditional distributions, following the LOMAR framework. Edge color represents directionality (red: negative influence; blue: positive influence); edge thickness indicates influence strength. Node size reflects centrality.

### 2.3 Simulated Forecasting for a Hypothetical Phantom Site

To evaluate Q-net’s capacity for forecasting microbial dynamics in data-sparse scenarios, we performed a simulation using a “phantom” wastewater site—an entirely synthetic subject initialized with randomized microbial abundances and no observed history. While this does not assess generalization to real-world unseen data, it illustrates the model’s generative capability to simulate coherent ecological dynamics under hypothetical conditions.

Figure 6 displays the model’s 30-week forecasts (weeks 60–90) for the top 10 most abundant taxa. These results demonstrate that even in the absence of subject-specific training data, Q-net can synthesize plausible, temporally coherent microbial trajectories—providing a potentially useful tool for exploratory analysis or ecological hypothesis generation. However, we emphasize that such simulations do not represent out-of-sample generalization performance and should be interpreted accordingly.

**Figure 6.**
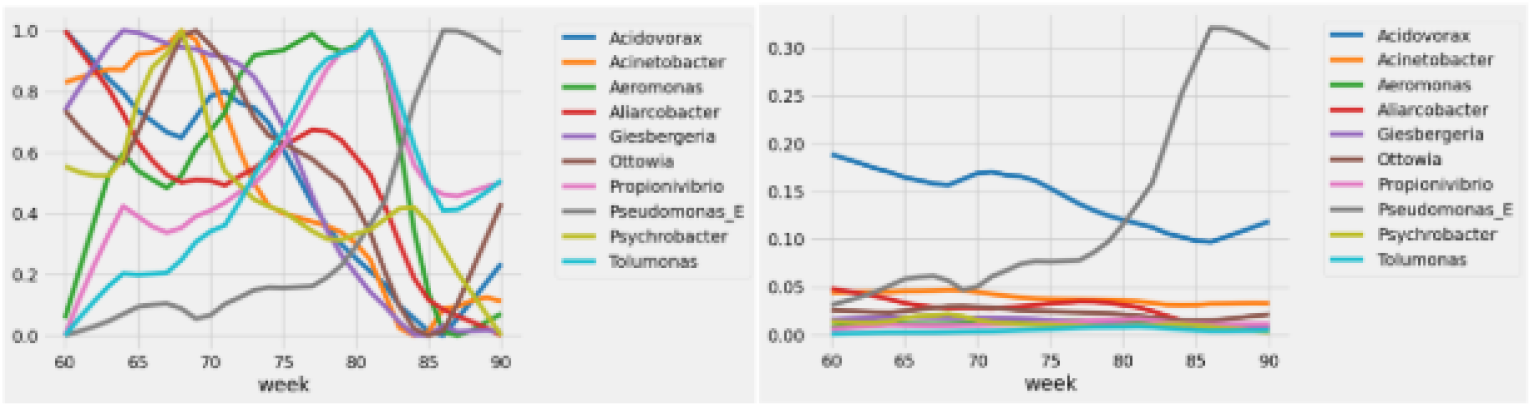
Predicted relative abundance time series for the top 10 most abundant taxa at a synthetic phantom site generated using Q-net. The left panel shows normalized abundances (scaled to [0,1]) to highlight dynamic trends across taxa, while the right panel presents unnormalized relative abundances, preserving original magnitude differences.

**Figure 7.**
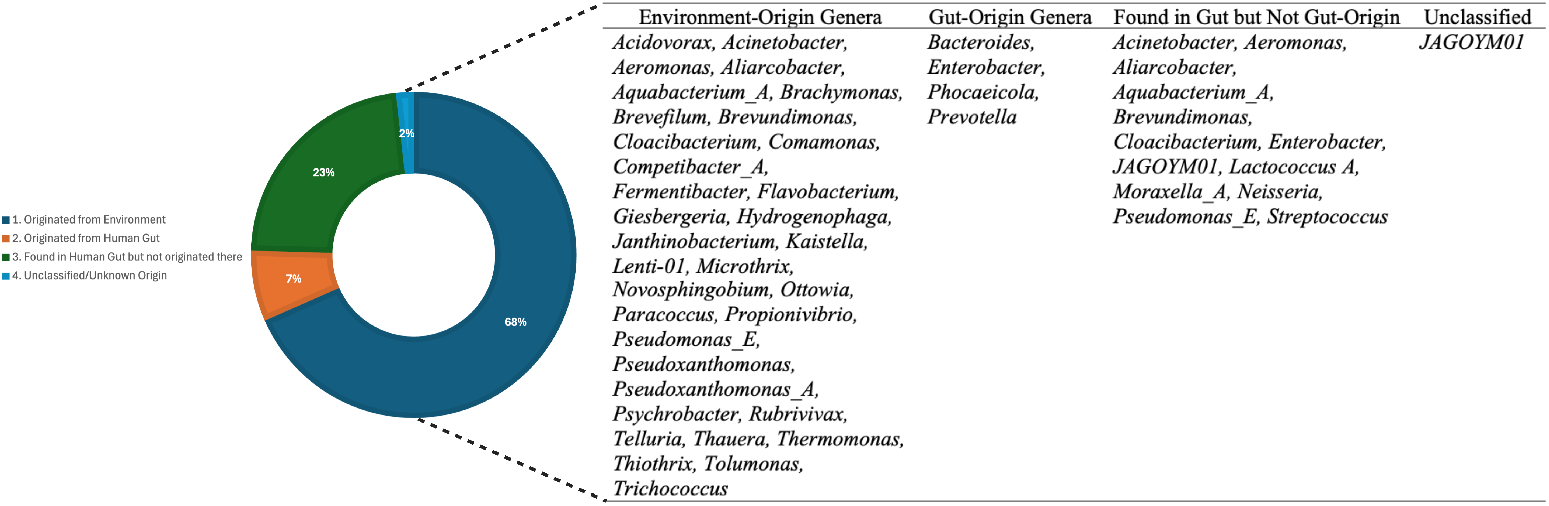
Donut chart illustrating the distribution of bacteria genera based on their origin. Table on right further breaks down the genera within each category, listing specific bacterial names.

**Figure 8.**
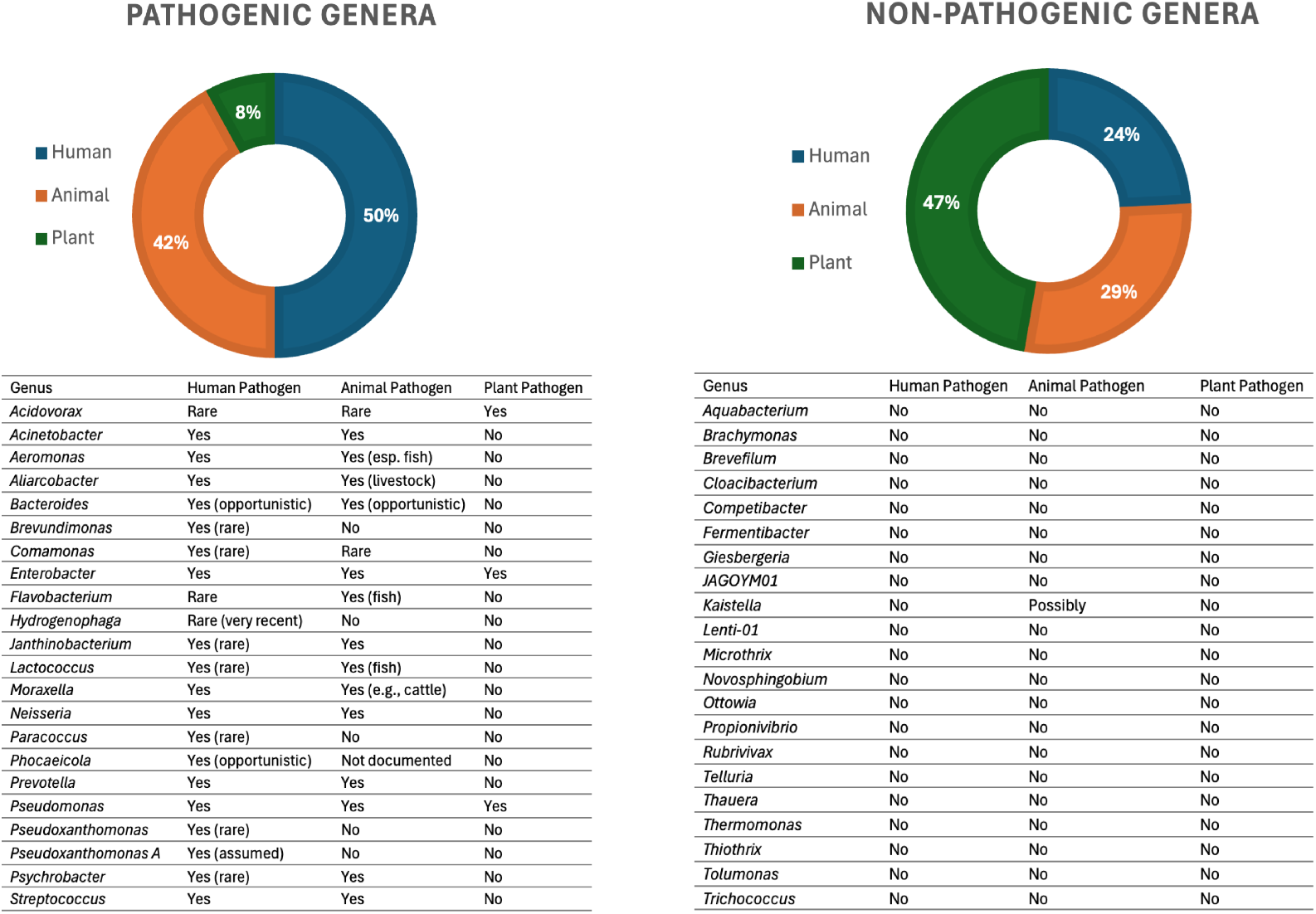
The top portion features two donut charts illustrating the percentage distribution of each host type for both pathogenic (left) and non-pathogenic (right) genera. Below the charts, two detailed tables list individual genera and indicate whether they are human, animal, or plant pathogens, often specifying their prevalence.

**Figure 9.**
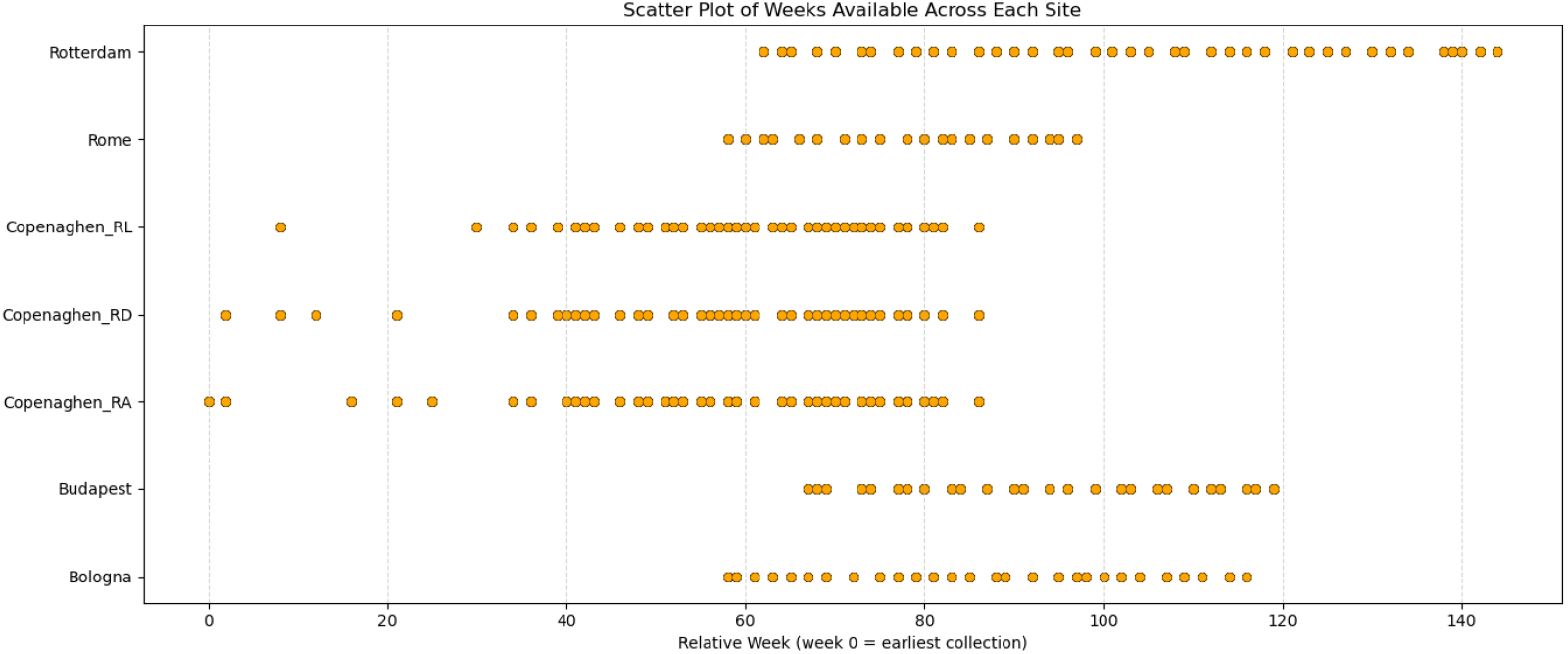
Scatter plot showing available sampling weeks across different wastewater treatment sites. Each point represents one metagenomic sample collected at the corresponding site and week. The x-axis indicates relative week (week 0 = earliest sample collected).

### 2.4 Source Attribution of Microbial Communities

Wastewater surveillance relies heavily on the ability to discern the origin of microbial signals. As demonstrated by Bescei et al., quantifying human-specific markers like crAssphage (a highly abundant human gut bacteriophage) and human mitochondrial DNA offers a valuable approach to differentiate between human gut microbes and environmental bacteria. Building on this, our analysis further categorized bacterial genera with abundances greater than 0.5% by their source. The results, visually represented in Figure 10, clearly show that a substantial proportion, approximately 70%, of the identified bacterial genera originated from the environment, indicating a dominant environmental microbial contribution in the wastewater samples. Beyond source attribution, we extended our classification to assess the pathogenic potential of these microbes, as shown in Fiugre **??**. By comparing the MAGs to established literature on microbial pathogenicity, we were able to determine whether the identified bacteria were pathogenic. This detailed classification revealed that among all pathogenic bacteria detected, a significant 50% were found to be human pathogens. Conversely, only a minor fraction of the identified bacteria posed a pathogenic threat to plants. This comprehensive approach, combining source tracking with pathogenicity assessment, provides a more holistic understanding of the microbial landscape in wastewater and its potential implications for public and environmental health.

**Figure 10.**
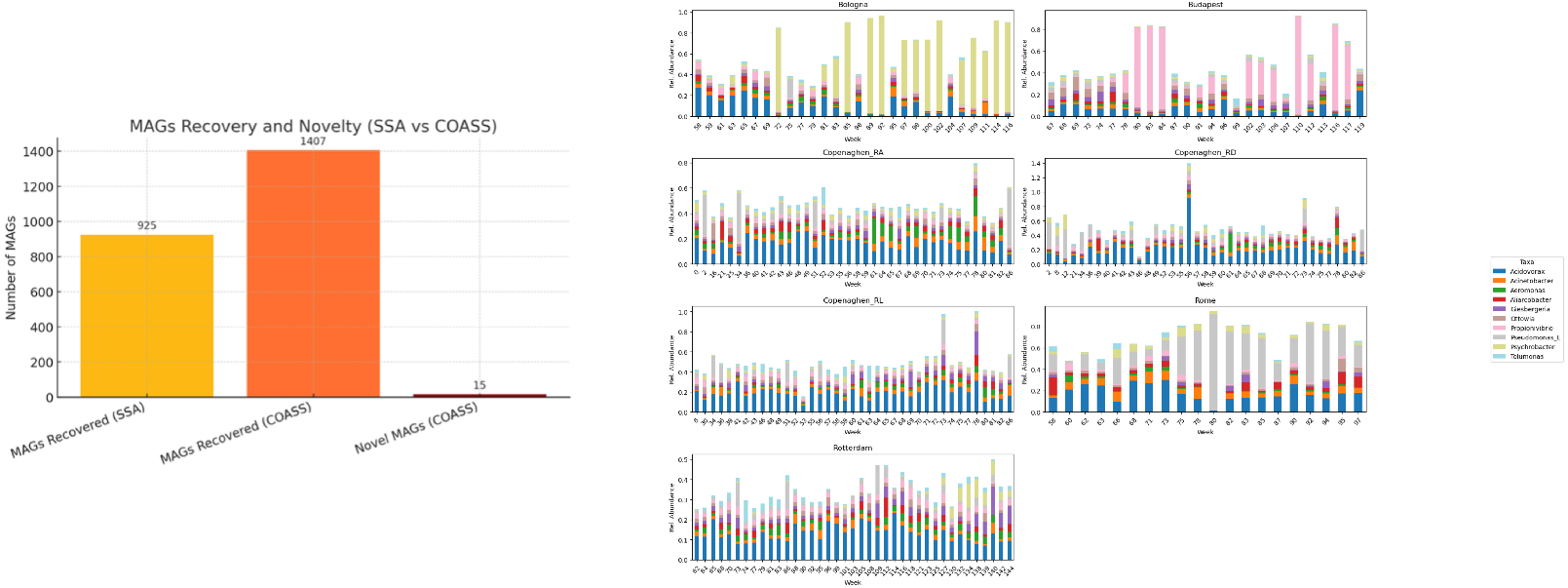
Comparison of MAG recovery and bacterial community dynamics. (Left) Comparison of MAG recovery using SSA and COASS methods, highlighting the enhanced recovery and novelty achieved by COASS. Data adapted from Bescei et al. (Right) Temporal dynamics of the top 10 most abundant bacterial genera across seven wastewater treatment plants in five European cities.

## 3 Discussion

The escalating global connectivity and the persistent threat of emerging infectious diseases underscore the urgent need for a paradigm shift in public health surveillance. WBE has emerged as a promising, non-invasive, and cost-effective tool, providing a macroscopic lens into population health dynamics. Our study significantly advances the utility of WBE by introducing a novel digital twin framework for urban wastewater microbiomes, moving beyond descriptive analyses to achieve predictive intelligence and source-attributed surveillance of pathogens.

Most MAGs belong to novel species, which help us better understand microbial populations and their interaction with the environment they liveSetubal2021. The type of assembly strategies used to reconstruct the microbial genomes directly affects the MAGs. Through the implementation of COASS strategies, MAG’s recovery was significantly improved over SSA. Notably, COASS uniquely identified novel microbial linkages, revealing previously uncharacterized taxa. The temporal dynamics observed across diverse European wastewater treatment facilities further highlight spatial heterogeneity and time-dependent fluctuations, shaped by environmental conditions, operational factors, and biofilm-associated shedding events.

The core contribution of our work lies in adapting and applying the Q-net digital twin framework to time series sewage dataset to forecast microbial abundance. Trained on longitudinal metagenomic data, Q-net demonstrated exceptional predictive performance in forecasting microbial abundance trajectories. High R^2^ values (exceeding 0.97 for many taxa) and a near-perfect R^2^ of 0.998 at the final forecast horizon attest to the model’s ability to capture both global and taxon-specific dynamics. This marks a significant shift from static, correlative studies to a truly predictive epidemiological framework. Beyond its forecasting accuracy, Q-net provides a critical advantage in model interpretability. By leveraging the CITs to model individual taxa, the framework offers transparent insights into ecological dependencies and temporal lag effects. For instance, the CIT for Enterobacter_65 revealed strong predictive relationships with past abundance states of other taxa such as *Neisseria_67*. Moreover, the directional influence network inferred from Q-net’s marginalization procedures further elucidates microbial relationships, identifying key regulatory taxa that exert influence over community structure. Beyond accurate forecasting, Q-net offers a distinct advantage through its interpretability. The use of CITs to model each microbial taxon provides a transparent view into the complex ecological dependencies and temporal lag effects within the microbiome. As illustrated by the example CIT for *Enterobacter_65*, the model can identify strong predictors of future dynamics based on the past abundance states of other taxa (e.g., Neisseria_67). This interpretability is crucial for uncovering underlying microbial interactions and for elucidating the mechanisms driving microbiome shifts and pathogen dynamics, a feature often lacking in more opaque machine learning models. The inferred directional influence network, derived from Q-net’s marginalization procedures, further illuminates these relationships, highlighting key regulatory taxa that may exert significant control over community structure.

Crucially, the digital twin enables source attribution, effectively distinguishing and quantifying contributions from human versus environmental (biofilm-associated) sources. This represents a pivotal advancement for targeted public health interventions, facilitating more accurate identification of contamination origins. Additionally, the framework’s ability to predict the emergence and prevalence of specific pathogens provides an invaluable early warning system.

## 4 Materials and Methods

### 4.1 Study Design and Data Source

We performed a computational reanalysis of a previously published longitudinal metagenomic dataset from^17^, which profiled the microbial dynamics of untreated sewage across seven treatment plants in five European cities over a two-year period. The original study applied shotgun metagenomic sequencing to characterize microbial community structure, antimicrobial resistance (AMR) gene content, and seasonal variability. Sample collection, DNA extraction, library preparation, sequencing, and metagenomic assembly procedures are fully described in the original publication and its supplementary materials. Sampling coverage varied across sites and time, with detailed weekly availability shown in Figure 9

### 4.2 MAG Recovery and AMR Quantification (As Reported in Source Study)

Briefly, the authors assembled over 25 million contigs via both single-sample assembly (SAA) and co-assembly (COASS) approaches. MAGs were binned using k-mer and depth-based methods, resulting in 2,332 medium-to high-quality MAGs, including 1,334 genomes from previously undescribed species. AMR genes were annotated using Resfams and ResFinder databases as described in^17^. As illustrated in Figure 10, the implementation of COASS strategies significantly enhanced the recovery of MAGs compared to SSA. Specifically, COASS yielded 1407 species’ genomes, substantially expanding upon the 925 species’ genomes recovered via SSA. Furthermore, the co-assembly process was particularly effective in uncovering novel microbial lineages, with 15 of the 19 unclassified species exclusively obtained through this method, underscoring its capacity to reveal previously undescribed taxa from sewage.

Similarly, the temporal dynamics of the top 10 abundant bacterial genera were observed across seven wastewater treatment facilities located in five major European cities, Bologna, Budapest, Rome, Rotterdam, and three sites in Copenhagen (RA, RD, RL), over multiple weeks. Each bar represents the relative abundance of bacterial taxa per week, highlighting both spatial heterogeneity and temporal fluctuations in community composition. Notably, *Pseudomonas_E* and *Acidovorax* frequently dominate across several locations, particularly in Rome and Rotterdam, while Budapest exhibits periodic spikes in *Psychrobacter* abundance. The microbial communities in Copenhagen sites show more balanced distributions, though transient blooms, such as a peak in *Cloacibacterium* in RD, are evident. These trends underscore the dynamic and site-specific nature of wastewater microbial ecosystems, potentially influenced by localized environmental conditions, operational differences in treatment infrastructure, or biofilm-associated shedding events.

To quantify AMR genes, the researchers mapped trimmed sequencing reads from all libraries using kma (v1.2.8) with specific flags (−cge and −lt1) against the ResFinder database. The ResFinder database is a curated collection of antimicrobial resistance sequences. Settings for kma allowed only one query sequence per template with default penalty values. The aligned sequence fragments were then summed to representative sequences, which were grouped by similar ResFinder alleles using 90% identity and 90% coverage thresholds with usearch (v11.0.667) cluster_fast.

### 4.3 Modeling Temporal Microbial Dynamics Using Q-net

To model the time-dependent structure of microbial communities in longitudinal sewage metagenomic data, we adopted the Q-net framework introduced by^15^. Q-net constructs a generative model, referred to as a digital twin, that captures high-order temporal dependencies among microbial taxa without relying on pre-specified interaction networks. This approach enables predictive simulation of microbial ecosystem trajectories from sparse early-time observations.

In our analysis, we focused on the subset of data corresponding to weeks 60 to 90 (relative to the earliest sample collected), as this interval provided the most consistent sampling coverage across sites (see Figure 9). Timepoints outside this window were excluded due to inconsistent temporal resolution or site-specific gaps.

### 4.4 Quantization of Relative Abundances

Let *x*_*i,t*_ ∈ [0, 1] denote the relative abundance of microbial taxon *i* ∈ {1, …, *N*} at timepoint *t* ∈ {1, …, *T*}. To mitigate the effects of compositionality and to facilitate categorical modeling, we discretized the continuous abundance values into *Q* quantile-based bins using the empirical distribution of each taxon:

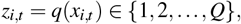

where *q*(·) maps abundance values to quantized bin levels. This transformation yields a structured categorical representation ℒ= {*z*_*i,t*_}, which serves as the input to Q-net.

### 4.5 Q-net Model Architecture

Q-net is formulated as a structured ensemble of conditional inference trees {*T*_*i,t*_}, where each tree models the conditional distribution of a specific taxon–timepoint pair:

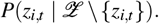

Each decision tree *T*_*i,t*_ is trained to predict the state of *z*_*i,t*_ using the remaining variables ℒ \{*z*_*i,t*_} as features. No temporal locality constraints are imposed, allowing the model to capture long-range dependencies across taxa and time. Node splits within each tree are selected using permutation-based statistical tests, which limit overfitting by enforcing significance thresholds.

The resulting model defines a joint distribution over the full trajectory space ℒ ^*N×T*^ through the product of conditional estimators:

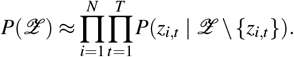

Because the inference procedure is stochastic, we generate an ensemble of Q-nets through repeated bootstrapped training to ensure robustness.

### Model Evaluation

Forecasting performance was assessed by conditioning Q-net on early observations and predicting microbial abundances at later timepoints. Let 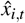 denote the predicted abundance for taxon *i* at time *t*, and *x*_*i,t*_ the observed value. Model accuracy was quantified using the coefficient of determination *R*^2^:

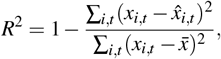

where 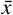 denotes the mean abundance across all samples. We report ensemble-averaged *R*^2^ scores across bootstrap replicates to evaluate forecasting capability and assess generalizability across timepoints and study sites.

## 5 Limitation and Future Work

This study has several limitations that should be considered when interpreting the findings. First, the analysis relies entirely on a publicly available dataset from^17^, meaning we had no control over sampling protocols, environmental variables, or sequencing consistency, which may influence the robustness of the forecasting outputs. Second, although the original dataset includes annotations for source attribution and AMR, our work did not introduce or evaluate new methods for these aspects. Finally, the model does not incorporate potentially relevant environmental covariates such as temperature, precipitation, or demographic factors, which may improve forecasting accuracy and ecological insight in future iterations.

Building on this foundational digital twin framework, future work will focus on expanding its temporal and spatial scope to improve generalizability. Incorporating longer time series from diverse geographic regions and varying environmental conditions, along with detailed metadata such as temperature, precipitation, industrial discharges, and demographic trends, will enable more robust and context-aware predictions. Integrating additional omics layers, such as metatranscriptomics and metaproteomics, will also offer a dynamic view of microbial function, allowing us to move beyond genetic potential and directly observe active expression of AMR genes, virulence factors, and biofilm-associated processes.

To enhance usability and impact, we aim to incorporate “what-if” simulation capabilities, allowing users to test hypothetical scenarios such as the introduction of novel pathogens or AMR genes and evaluate the effects of public health interventions or operational changes. This will be supported by the development of an interactive 3D platform, offering an intuitive visualization of microbial dynamics, source contributions, and forecast trajectories. Coupling the digital twin with real-time data streams and operational infrastructure models will further enable wastewater treatment operators and public health authorities to make proactive, data-driven decisions for surveillance, outbreak response, and treatment optimization. In parallel, we aim to refine the model’s source attribution capabilities by introducing molecular markers to differentiate between human, environmental, and industrial inputs, supported by targeted validation studies. Applications include optimizing treatment parameters, predicting biofilm development, and enabling early intervention based on pathogen or AMR gene emergence. Ultimately, close collaboration with public health and wastewater stakeholders will be essential to validate and translate the model’s predictions into actionable interventions, bridging the gap between environmental surveillance and public health response.

## Acknowledgements

The authors gratefully acknowledge the support from the National Science Foundation RII Track-2 award 1920954 and Track-1 award1849206, Institutional Development Award (IDeA) from the National Institute of General Medical Sciences of the National Institutes of Health (P20GM103443)

## Author contributions statement

Conceptualization, B.D.S.G. and E.Z.G.; methodology, B.D.S.G., E.Z.G., S.A., and T.D.; software, B.D.S.G.; validation, D.B. and R.Y.; formal analysis, E.Z.G.; data curation, M.R., S.A., and T.D.; writing—original draft preparation, M.R.; writing—review and editing, B.D.S.G., M.R., and N.M.; visualization, B.D.S.G. and N.M.; supervision, E.Z.G.

## References

1. Buckee, C. O. et al. Productive disruption: opportunities and challenges for innovation in infectious disease surveillance. BMJ Glob. Heal. 3, e000538, DOI: 10.1136/bmjgh-2017-000538 (2018).

2. Chau, K. et al. Systematic review of wastewater surveillance of antimicrobial resistance in human populations. Environ. Int. 162, 107171, DOI: 10.1016/j.envint.2022.107171 (2022).

3. Gagliano, E., Biondi, D. & Roccaro, P. Wastewater-based epidemiology approach: The learning lessons from covid-19 pandemic and the development of novel guidelines for future pandemics. Chemosphere 313, 137361, DOI: 10.1016/j.chemosphere.2022.137361 (2023).

4. Chen, C. et al. Wastewater-based epidemiology for covid-19 surveillance and beyond: A survey. Epidemics 49, 100793, DOI: 10.1016/j.epidem.2024.100793 (2024).

5. LaMartina, E. L., Mohaimani, A. A. & Newton, R. J. Urban wastewater bacterial communities assemble into seasonal steady states. Microbiome 9, DOI: 10.1186/s40168-021-01038-5 (2021).

6. Chahal, C. et al. Pathogen and Particle Associations in Wastewater, 63–119 (Elsevier, 2016).

7. Saini, S., Tewari, S., Dwivedi, J. & Sharma, V. Biofilm-mediated wastewater treatment: a comprehensive review. Mater. Adv. 4, 1415–1443, DOI: 10.1039/d2ma00945e (2023).

8. Nahum, Y., Muhvich, J., Morones-Ramirez, J. R., Casillas-Vega, N. G. & Zaman, M. H. Biofilms as potential reservoirs of antimicrobial resistance in vulnerable settings. Front. Public Heal. 13, DOI: 10.3389/fpubh.2025.1568463 (2025).

9. Faraway, J. et al. Challenges in realising the potential of wastewater-based epidemiology to quantitatively monitor and predict the spread of disease. J. Water Heal. 20, 1038–1050, DOI: 10.2166/wh.2022.020 (2022).

10. Zhu, W., Wang, D., Li, P., Deng, H. & Deng, Z. Advances in wastewater-based epidemiology for pandemic surveillance: Methodological frameworks and future perspectives. Microorganisms 13, 1169, DOI: 10.3390/microorganisms13051169 (2025).

11. Alsalloum, G. A., Al Sawaftah, N. M., Percival, K. M. & Husseini, G. A. Digital twins of biological systems: A narrative review. IEEE Open J. Eng. Medicine Biol. 5, 670–677, DOI: 10.1109/ojemb.2024.3426916 (2024).

12. Neupane, A. Advanced news aggregation and content generation using llms and nlp algorithms. Eur. J. Appl. Sci. Eng. Technol. (EJASET) 3, 45–50 (2025). Accessed: 2025-07-22.

13. Banjade, M. et al. Deep learning and gen ai based system for ingredient recognition and recipe insight. Int. J. Sci. Eng. Technol. 12, – (2024). Preprint available on ResearchGate.

14. Hothorn, T., Hornik, K. & Zeileis, A. Unbiased recursive partitioning: A conditional inference framework. J. Comput. Graph. Stat. 15, 651–674 (2006).

15. Sizemore, N. et al. A digital twin of the infant microbiome to predict neurodevelopmental deficits. Sci. Adv. 10, DOI: 10.1126/sciadv.adj0400 (2024).

16. Wu, J. & Koelzer, V. H. Towards generative digital twins in biomedical research. Comput. Struct. Biotechnol. J. 23, 3481–3488, DOI: 10.1016/j.csbj.2024.09.030 (2024).

17. Becsei, et al. Time-series sewage metagenomics distinguishes seasonal, human-derived and environmental microbial communities potentially allowing source-attributed surveillance. Nat. Commun. 15, DOI: 10.1038/s41467-024-51957-8 (2024).

18. Zhang, Y. et al. Microprophet: A digital twin framework for predicting microbial community dynamics with personalized precision. DOI: 10.1101/2025.05.08.652793 (2025).

19. Setubal, J. C. Metagenome-assembled genomes: concepts, analogies, and challenges. Biophys. Rev. 13, 905–909, DOI: 10.1007/s12551-021-00865-y (2021).

